# Isolation and characterization of a novel bacteriophage, Kapi1, capable of O-antigen modification in commensal *Escherichia coli*

**DOI:** 10.1101/2021.04.09.439263

**Authors:** Kat Pick, Tracy Raivio

**Affiliations:** Department of Biological Sciences, University of Alberta, Edmonton, Alberta, Canada

## Abstract

In this study, we describe the isolation and characterization of novel bacteriophage Kapi1 (vB_EcoP_Kapi1) isolated from a strain of commensal *Escherichia coli* inhabiting the gastrointestinal tract of healthy mice. We show that Kapi1 is a temperate phage integrated into tRNA *argW* of strain MP1 and describe its genome annotation and structure. Kapi1 shows limited homology to other characterized prophages but is most similar to the phages of *Shigella flexneri*, and clusters taxonomically with P22-like phages. Investigation of the lifestyle of Kapi1 shows that this phage displays unstable lysogeny and influences the growth of its host. The receptor for Kapi1 is the lipopolysaccharide O-antigen, and we further show that Kapi1 alters the structure of its hosts O-antigen in multiple ways. We hope to use MP1 and Kapi1 as a model system to explore molecular mechanisms of mammalian colonization by *E. coli* and ask what the role(s) of prophages in this context might be.

**Importance:** Although research exploring the microbiome has exploded in recent years, our understanding of the viral component of the microbiome is lagging far behind our understanding of the bacterial component. The vast majority of intestinal bacteria carry prophages integrated into their chromosomes, but most of these bacteriophages remain uncharacterized and unexplored. Here, we isolate and characterize a novel temperate bacteriophage infecting a commensal strain of *Escherichia coli.* We aim to explore the interactions between bacteriophages and their hosts in the context of the gastrointestinal tract, asking what role(s) temperate bacteriophage may play in growth and survival of bacteria in the gastrointestinal tract. Understanding the fundamental biology of commensal bacteria in the gastrointestinal tract can inform the development of novel antimicrobial or probiotic strategies for intestinal infections.

## Introduction

*Escherichia coli* is a Gram-negative bacterium normally inhabiting the lower gastrointestinal tract of humans and other warm-blooded animals (1). Despite being one of the most widely studied prokaryotic model organisms, there remains an immense complexity to the lifestyle of *E. coli* that we are only beginning to appreciate; one of these layers of complexity is the interactions between *E. coli* and the bacteriophages that infect it. Bacteriophage (or simply phage) exhibit two main lifestyles; lytic phage infect and immediately begin replicating within their host, eventually causing cell lysis and releasing progeny phages. Temperate phage integrate into the genome of their host where they reside as mostly dormant prophages, replicating with the host chromosome and being disseminated into daughter cells. Once the host cell experiences stress such as DNA damage or starvation, the prophage excises from the chromosome and enters the lytic cycle to ensure its’ own survival (2). Temperate phage have been gaining attention as we begin to appreciate their abundance; it has been estimated that approximately half of all sequenced bacterial genomes contain intact prophage, and even more contain prophage elements (3). Interestingly, the abundance of temperate phage residing in the commensal gut microbiome of mice appears to be even higher (4), indicating that temperate phage may play a role in bacterial community dynamics during colonization. Indeed, many recent studies and reviews have highlighted the importance of phages in the microbiome community (5–8).

One of the ways in which temperate phage influence the biology of their hosts is through lysogenic conversion. During lysogenic conversion, “extra” genes, called morons, encoded on the prophage are expressed in the host cell during lysogeny. These morons influence the biology of the host cell, without affecting the phage life cycle. One form of lysogenic conversion is seroconversion, in which bacteriophages encode proteins that alter the structure of the host lipopolysaccharide (LPS) O-antigen. The most well-known seroconverting phage are those that infect *Shigella flexneri*; lysogeny with these phage results in modification of the O-antigen through either glucosylation or O-acetylation, leading to a change in bacterial serotype (9). This can have different benefits for the bacterial host including antigenic variation and immune evasion, since the mammalian innate immune system mounts a serotype-specific antibody response (9). Beyond immunogenicity, LPS is an essential component of the outer membrane that is important for membrane stability and barrier function (10).

Here, we describe the isolation and characterization of novel bacteriophage Kapi1, capable of O-antigen modification. Kapi1 was isolated from a wild commensal strain of *E. coli*, and the phage genome was sequenced and compared to other characterized phages. We also report the identification of the O-antigen as the receptor for Kapi1 and show that this phage displays an unstable temperate lifestyle. Our characterization of Kapi1 suggests that it has a significant impact on the physiology and lifestyle of *E. coli*; it should serve as an excellent model system to explore the impact of bacteriophage on *E. coli* colonization of the mammalian gastrointestinal tract.

## Results and Discussion

### Kapi1 is a novel *Podoviridae* with a narrow host range

Recently, Lasaro et al. (2014) showed that the Cpx, Arc, and Rcs two-component systems found in *Escherichia coli* were required for colonization of the murine gastrointestinal tract by a strain of commensal *E. coli*, MP1. We began performing competitions between Cpx, Arc, and Rcs mutants and WT MP1 *in vitro* to further explore the molecular mechanisms behind the observed colonization phenotypes. MP1, MP7, and MP13 are identical strains except for the presence of fluorescent plasmids pML8 and pAS07 integrated into the chromosomes of MP7 and MP13 respectively, at the λ attachment site (11). Because MP7 and MP13 are marked with *mcherry* and *gfpmut3.1*, these strains are easily distinguishable during co-culture competition experiments. Unexpectedly, when co-culturing MP13 *rcsB* mutants with MP7, we found that the *rcsB* mutants strongly outcompeted the WT (data not shown). Because Lasaro et al (2014) showed that mutation of *rcsB* decreased competitiveness in a mouse colonization model, we wondered if this reflected a differential ability of the Rcs mutant to compete *in vitro* vs *in vivo* and set out to investigate this. Because of the strong competitive advantage, we hypothesized that the *rcsB* mutant could perhaps be directly killing the WT in some way. To test if there was a bactericidal factor secreted by the *rcsB* mutant, we isolated the supernatant from cultures of MP13 *rcsB* mutants and spotted it onto lawns of MP7. Unexpectedly, the supernatant cleared the MP7 lawn, and serial dilutions of the supernatant showed spotty clearing, reminiscent of phage plaques. We then screened our entire strain collection of all strains derived from MP1, MP7, and MP13 for their abilities to produce plaques on each other. A clear trend emerged; the supernatants of MP1 and MP13 background strains could produce plaques on lawns of MP7, but supernatant derived from MP7 background strains could not produce plaques on either MP1 or MP13. Thus, we began identification and characterization of the phage found in MP1 and MP13.

Because our cultures of MP1 and MP13 containing phage appeared healthy, we hypothesized that the phage in these cultures may be lysogenic, as a lytic phage would be more likely to lyse the cultures resulting in a visible reduction in cell density and poor growth. Analysis of the previously published genome sequence for MP1 (11) using the prophage identification tool PHASTER (12, 13) revealed six putative prophages integrated into the chromosome of MP1 (Figure 1A). Of these, only one prophage was scored as intact by PHASTER (12, 13) (Figure 1A); we hypothesized that this prophage was the most likely candidate for the phage plaques we had observed because of the completeness of the prophage sequence. To confirm this, we performed PCR on colonies of MP1, MP7 and MP13, as well as phage lysates prepared from MP1 and MP13 with three primer pairs targeting the coat, portal, and tail proteins within the Intact_1 prophage region identified by PHASTER. Bands were consistently observed for MP1 and MP13 colonies and phage lysates and were consistently absent for MP7 colonies for all primer pairs (data not shown). This indicated that the phage present in MP1 and MP13 cultures likely corresponds to the Sf101-like Intact_1 prophage region identified by PHASTER in the MP1 genome. Despite several attempts to induce phage from MP7 using either DNA damaging agents or cellular starvation, phage could never be isolated from MP7. Since Lasaro et al. (2014) previously showed that MP7 and MP13 had equal competitive indices *in vivo*, we decided to investigate this further. Upon testing the original stock of MP7 isolated by Lasaro et al. (2014), we found that this strain does in fact contain the Intact_1 prophage region (data not shown), and that only our stock of MP7 lacks the Intact_1 prophage region. These findings are important as they demonstrate that any competitive advantage that would have been provided by carrying the Intact_1 prophage did not play a role in the findings of Lasaro et al. (2014), since both MP7 and MP13 contain the prophage. We propose that our stock of MP7 which lacks the Intact_1 prophage region should be renamed to KP7, in order to avoid confusion with the original MP7 stock which does contain the Intact_1 prophage region.

**Figure 1.**
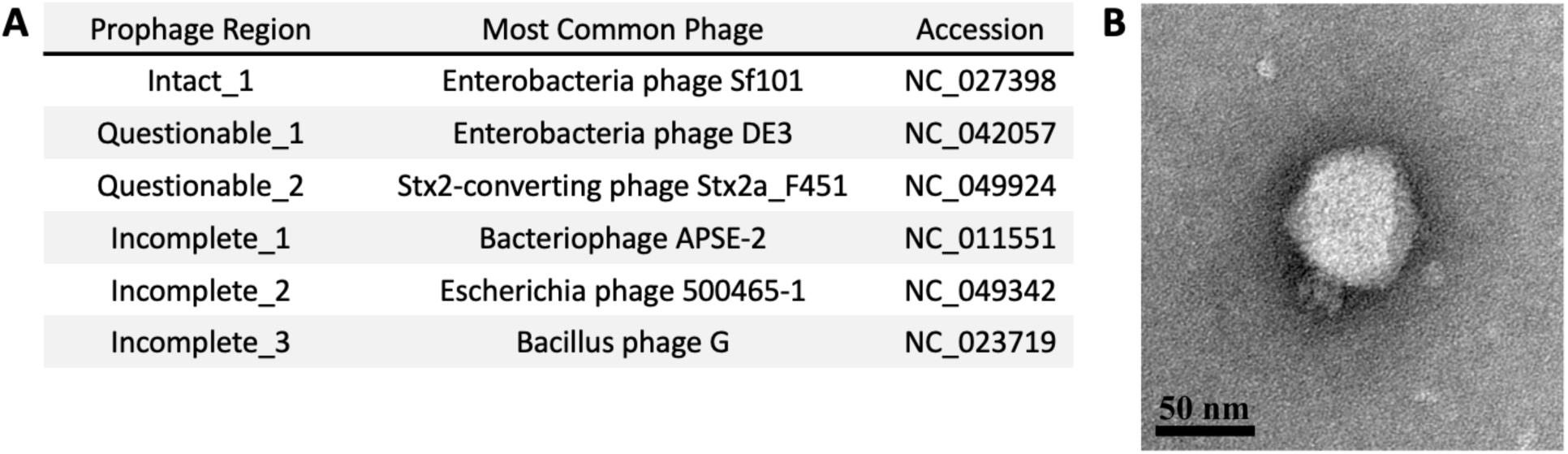
PHASTER analysis of MP1 and transmission electron microscopy image of Kapi1. (A) The genome of MP1 (accession JEMI01000000) (11) was analyzed for putative prophage regions using PHASTER (12, 13). Prophage regions identified by PHASTER are shown, including the most common phage from the NCBI viral database, and their corresponding accession numbers. Intact, questionable, and incomplete scores were assigned by PHASTER. (B) Kapi1 phage lysate was stained with 4% uranyl acetate on a copper grid, and viewed by transmission electron microscopy at 140,000x magnification.

Transmission electron microscopy (Figure 1B) of phage lysates collected from MP1 and MP13 revealed phage particles with a mean capsid diameter of 70.10 ± 2.92 nm and tail length 15.37 ± 1.45 nm placing this phage in the family *Podoviridae* and order *Caudovirale.* We screened 11 strains of *E. coli* (Top10, MG1655, TJ-LM, TJ-WM, TJ-LR, MC4100, W3110, BW25113, J96, E2348/69, Nissile 1917) and *Citrobacter rodentium* DBS100 for susceptibility to the phage (see Table S1). All strains were completely resistant to infection; from our strain collection, KP7 is the only strain that this phage can infect. This preliminary analysis suggests that the newly isolated phage has a relatively narrow host range. This phage forms diffuse plaques on KP7 with an average plaque diameter of 2.0 ± 0.22 mm after overnight incubation at 37 °C. Although the morphology of phage particles on TEM and plaques on soft agar overlays was consistent, to confirm that the Intact_1 prophage region is the only prophage in MP1 capable of active excision and lytic replication, we performed PCR on DNase-treated phage lysates using primers corresponding to each putative prophage region identified by PHASTER, and a *nuoA* primer pair to control for genomic DNA contamination. No bands were observed in KP7 lysates, and only the band corresponding to Intact_1 prophage was observed in the MP13 lysate (Figure S1).

### Kapi1 lacks sequence homology with other *Podoviridae*, and has a modular lambdoid genome

Although the whole genome sequence for MP1 has already been published (11), to be thorough and ensure that the isolated phage was truly a prophage and not introduced by contamination from our laboratory, whole genome sequencing was performed on our stocks of MP1, KP7, and MP13. We also aimed to find the integration site and characterize the genome of the isolated phage. As anticipated, the 39 kb Sf101-like region was present in the genomes of MP1 and MP13, and absent from the genome of KP7. Unfortunately, the phage genome was assembled into its own linear contig, not showing where it may be integrated into the host chromosome. This was observed in both the original MP1 sequence (11) and in our resequencing attempt; it is likely that the phage genome was in its circular form (ie. excised from the host chromosome) since we extracted DNA from late stationary phase cultures. Upon closer analysis of the original MP1 sequence (11), it appears that the ends of contig NZ_JEMI01000030, corresponding to the Intact_1 prophage region, are actually terminal repeats indicating that the sequence is in fact circular. To confirm this circularity, primers were designed pointing outward from each end of the phage contig (Figure 2A), and PCR and Sanger sequencing was performed on DNA extracted from phage lysates. The sequence of the PCR product obtained was consistent with the conclusion that the phage exists in a circular form at some point during its lifecycle and confirmed complete sequencing of the entire phage genome (Figure 2A).

**Figure 2.**
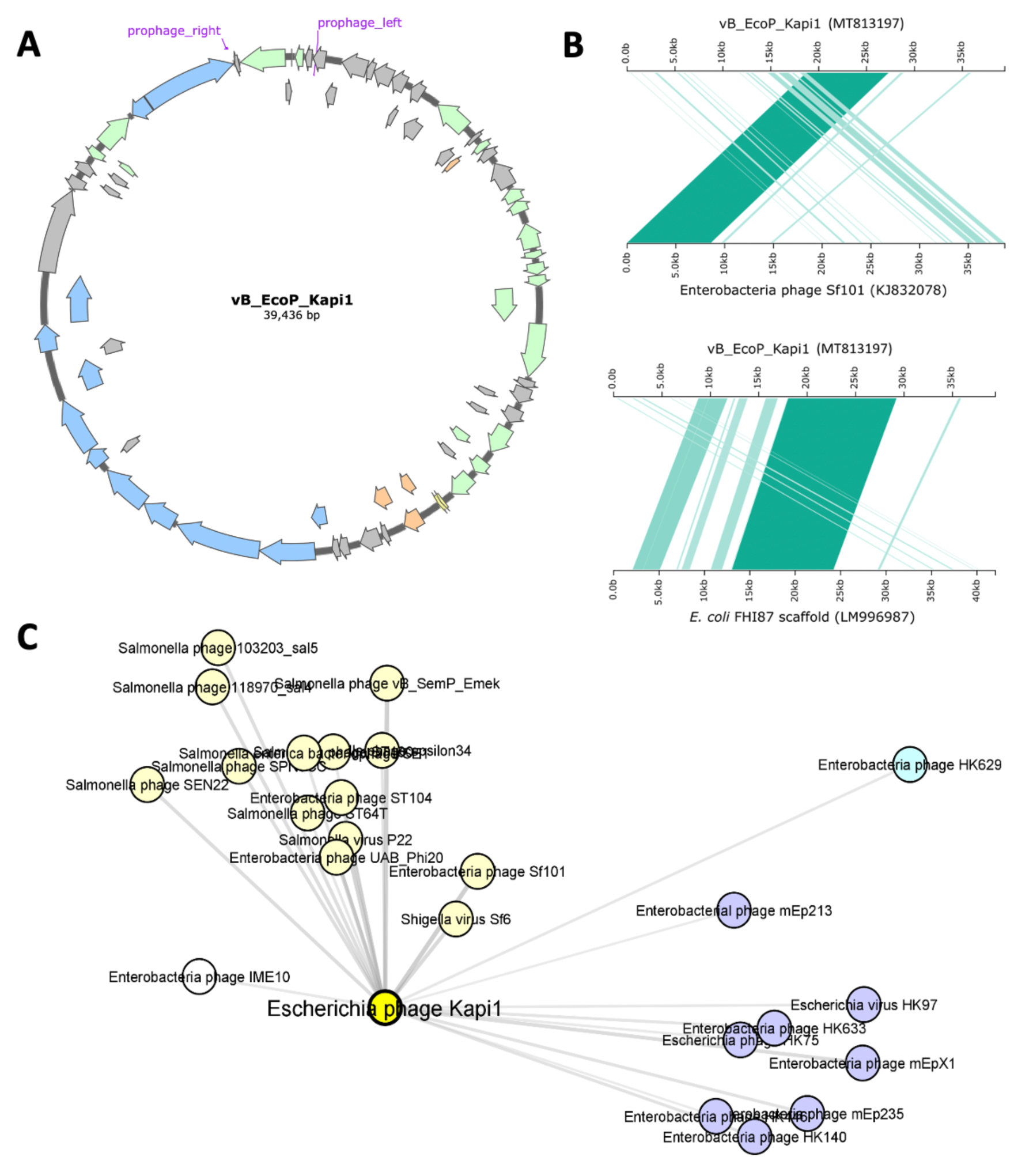
Genome of Kapi1. (A) Layout of the Kapi1 genome, with ORFs color coded as follows: lysis module in orange, structural/morphogenesis in blue, DNA replication/repair/regulators in green, tRNAs in yellow, and hypothetical proteins and proteins of unknown function in grey. Prophage_left and _right primers used to verify genome circularity are indicated in purple. (B) Alignment of Kapi1 genome with top NCBI BLASTn hits. The genome of vB_EcoP_Kapi1 (accession MT813197) was searched against the NCBI Blast nucleotide database, and the top hits were visualized with Kablammo (71); darker lines represent higher sequence homology. The upper panel shows the alignment of Kapi1 with the top hit from NCBI database without any filters, while the lower panel shows the top hit from NCBI database with the viral (taxid: 10239) filter applied. (C) vContact2 (17) clustering of Kapi1-related viruses, visualized with CytoScape (59). The first-neighbors network of Kapi1 was filtered to include only the top 50% strongest interactions with Kapi1, and with edges that do not directly connect to Kapi1 were removed for clearer visualization. An edge-weighted spring-embedded model was used so that the distance between viral nodes, and the darkness and thickness of edges connecting the nodes corresponds to the similarity between those viruses. Viral clusters are color-coded.

When the phage genome was analysed using BLASTn (14) with the viral filter (taxid: 10239), the top hit was Enterobacteria phage Sf101 (accession NC_027398.1) with 96.47% identify but only 33% query cover, indicating that this phage represents a novel viral species, with less than 95% nucleotide sequence similarity to any part of any other characterized phage (15). We thus named this novel phage vB_EcoP_Kapi1 (Kapi1, NCBI:txid2746235). When the viral filter is removed, the top hit still only shows 96.92% identity and 44% query cover to Kapi1 (*Escherichia coli* genome assembly FHI87, scaffold-10_contig-14.0_1_42036, accession LM996987.1). Visualization of the alignments between Kapi1 and these top BLASTn (14) hits (Figure 2B) showed that a ∼10 kb region of Kapi1 corresponding to the virion morphogenesis module is the most conserved region. Further, Kapi1 may represent a novel genus of the family *Podoviridae*, since it shares less than 50% nucleotide sequence similarity to any other characterized *Podoviridae* genus (15). Comparing the genome sequence of Kapi1 against the type-species for each *Podoviridae* genus in the ICTV 2019.v1 Master Species List (16), the top hit, Enterobacteria phage P22 (accession NC_002371.2), belonging to the genus *Lederbergvirus,* shares only 83.62% identity and 20% query coverage with Kapi1. The taxonomy of Kapi1 was further explored using vContact2 (17), this analysis showed that Kapi1 belongs to the same viral cluster as phages P22 (18) and Sf101 (19) (Figure 2C; Figure S2).

The genome of Kapi1 (accession MT813197) is 39,436 bp in length and represents 0.83% of the genome of MP1. The GC content of Kapi1 is 47.1%, slightly lower than the 50.6% of the host genome. Kapi1 has a modular genome structure typical of many lambdoid phages (Figure 2A) (20). Beginning from *xis*, the first region of the Kapi1 genome is rich in hypothetical proteins and proteins with unknown function. The next segment of the genome is characterized by the DNA replication/repair/regulation module; this region has a Lambda-like organization, with CIII, N, CI, cro, CII, O, P, and Q. This module is followed by two tRNAs immediately preceding the lysis module (holin, lysin, Rz). The next module is responsible for virion morphogenesis, encoding proteins responsible for the head assembly (terminases, scaffold, portal, coat), followed by tail assembly (DNA stabilization protein, tail needle knob, DNA transfer and ejection proteins). The final module is required for integration, including *xis, int,* and *attP.* For a detailed view of annotation and functional assignments for all CDS in Kapi1, see Table S3.

### Kapi1 integrates into the 3’ end of host tRNA *argW*

To begin our search for the integration site of Kapi1, we used BLASTn (14) to look for prophages similar to Kapi1 in the NCBI nucleotide database. We then analyzed the host chromosome surrounding these Kapi1-like prophages to find any similarities with the MP1 chromosome. We identified two contigs in MP1 whose ends shared significant similarity to the regions surrounding Kapi1-like prophages found in the NCBI database. The end of the first contig encodes the *dsdAXC* genes, while the end of the second contig encodes *yfdC, mlaA,* and *fadLIJ* genes. Primer pairs were designed to amplify the putative prophage-chromosome junctions; PCR products were then sequenced and aligned with the original sequences to determine the integration site of Kapi1, and its orientation in the host chromosome. Kapi1 integrates into the chromosome between genes *yfdC* and *dsdC*, with phage *int* gene closest to the chromosomal *yfdC* locus, and the phage *xis* gene on the opposite end of the prophage closest to the chromosomal *dsdC* gene (Figure 3A). Notably, the way in which the original MP1 sequence (11) was assembled actually captures the chromosome-prophage left_junction (Figure 3A), since the end of contig NZ_JEMI01000016 contains Kapi1 *xis* (annotated as *TorI* in the original sequence) directly downstream of *dsdC*. *yfdC* is a predicted inner membrane protein, belonging to the formate-nitrate transporter (FNT) family and may be involved in resistance to surfactants (21); *dsdC* is a transcriptional regulator involved in D-serine detoxification (22). This region of the genome is hypervariable among different *E. coli* pathotypes, and a variety of prophage and phage-like genes are often found at this locus (23).

**Figure 3.**
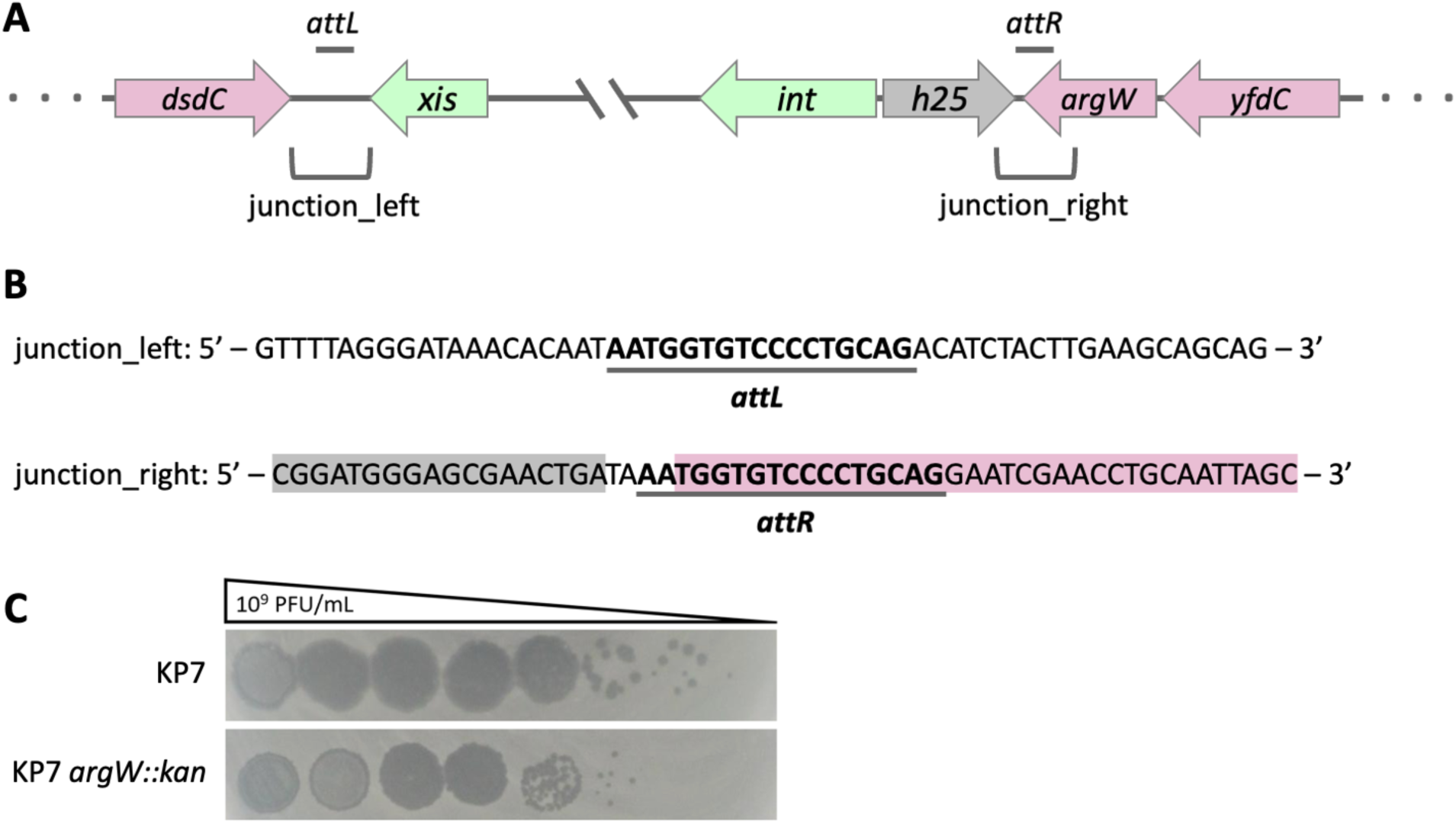
The integration site of Kapi1. (A) The structure of Kapi1 integrated into the host chromosome; host ORFs are indicated in pink, and phage ORFs are indicated in green and grey, in between the *attL* and *attR* sites. (B) Nucleotide sequences are provided for both the left and right host-prophage chromosome junctions, including 20 bp upstream and downstream of the bolded and underlined *att* sites. Grey shading on the junction_right sequence indicates the location of the Kapi1 *hyp25* gene, and pink shading indicates the host *argW* gene. (C) Kapi1 phage lysate was serially diluted and spotted onto soft-agar overlays of either WT KP7 or KP7 *argW::kan*.

With the integration site for Kapi1 identified, we then looked back at our whole-genome sequencing data for KP7, the strain lacking the Kapi1 prophage. In this strain, the integration locus was correctly assembled; the two contigs that we confirmed to surround the integrated Kapi1 prophage in MP1 were assembled into one complete contig in KP7. We noticed a tRNA-Arg in between *dsdC* and *yfdC* that was not annotated on the contigs surrounding the Kapi1 prophage in MP1. Since tRNAs are common integration sites for phage (24), this site was further investigated. When we investigated the chromosomes of MP1 and MP13 with the Kapi1 prophage integrated as described above, we noticed a 17 bp duplication on either end of the integrated prophage; this sequence (5’ – AATGGTGTCCCCTGCAG – 3’) is found at the 3’ end of the tRNA-Arg and is the putative Kapi1 *att* site (Figure 3B). To be clear, this tRNA-Arg is intact whether or not Kapi1 is integrated into the chromosome, since the 3’ end is maintained by the Kapi1 putative *attP* when it integrates into the chromosome. The *attB* site was not picked up by the auto-annotation programs in MP1 and MP13 since during sequencing the two contigs surrounding Kapi1 were not assembled into the correct scaffold, as they were in KP7.

Interestingly, the Kapi1 putative *attP* is identical to prophages Sf6 (25), HK620 (20), and KplE1 (26) except for the 5’ A which is excluded from the Sf6, HK620, and KplE1 *attP* sites. Like these phages, the Kapi1 *attP* lies between the *int* and *xis* genes, so upon integration into the host chromosome, the *int* and *xis* genes are located on either end of the prophage (Figure 3A). We generated a KP7 *argW::kan* mutant, in this mutant the *attB* core sequence is still intact, but the insertion of the kanamycin resistance cassette disrupts the sequence adjacent to the *attB* core, which may be important for recognition. As expected, this mutant can be infected by Kapi1, although at a reduced frequency when compared to KP7, and produces smaller plaques with an average diameter of 0.89 ± 0.25 mm (Figure 3C). Further experiments showed that KP7 *argW::kan* is still able to be lysogenized by Kapi1, but at a much lower frequency (13.5%) than the wildtype KP7 (100%). Notably, KP7 *argW::kan* lysogens retain the same level of resistance to kanamycin as their non-lysogenic counterparts. The observed reduced rate of lysogeny could be due to a reduced frequency of recognition of the intact *attB* core due to the adjacent sequence disruption by the kanamycin cassette, or possibly integration into other chromosomal sites (27); precise deletion of the putative *attB* core and further experimentation will help to clarify these results.

### The lifestyle of Kapi1 is temperate and shows unstable lysogeny

To investigate the lifestyle of Kapi1, we performed phage liberation assays. After only 24 hours of growth, ∼1×10^8^ PFU/mL can be isolated from standard laboratory cultures of lysogens (data not shown). The ratio of phage per cell (PFU/CFU) is only 0.030 ± 0.010 after 24 hours of growth and rises to 25.80 ± 7.23 PFU/CFU when identical cultures are grown with sub-inhibitory concentrations of mitomycin C (Figure 4A). The mitomycin C assay indicates that Kapi1 can be induced through the traditional SOS pathway in response to DNA damaging agents, although there appears to be a basal level of spontaneous induction in the absence of mitomycin C.

**Figure 4.**
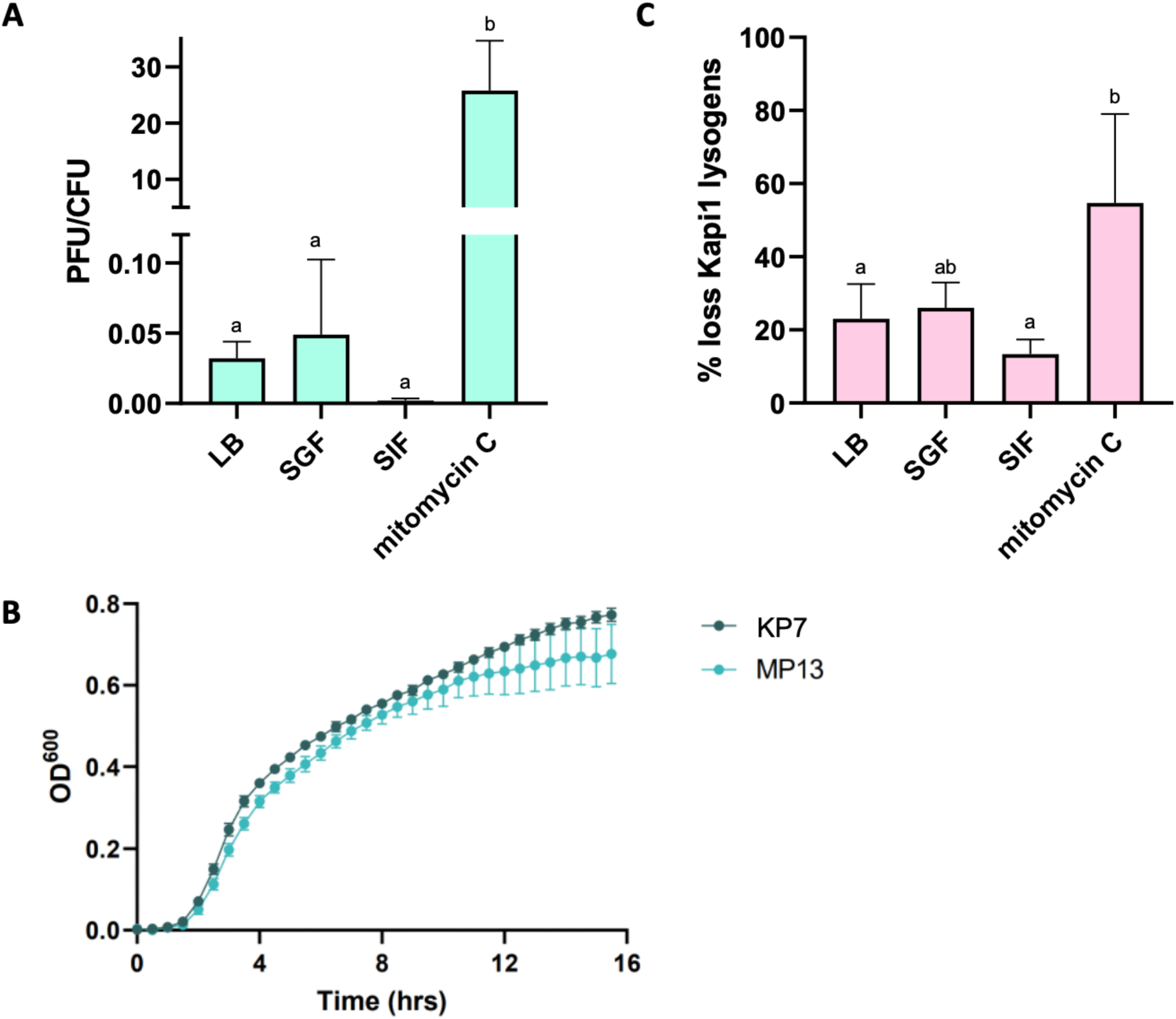
The lifestyle of Kapi1. (A) prophage liberation assay; cultures of MP13 (Kapi1 lysogen) were grown in LB, 50% LB 50% simulated gastric fluid (SGF) (29), 50% LB 50% simulated intestinal fluid (SIF) (29), or LB supplemented with 0.5 ng/uL mitomycin C for 24 hours. After 24 hours the number of cells were enumerated by spotting on LB plates, and the number of phages were enumerated by spotting on soft agar overlays prepared with KP121. PFU/mL was divided by CFU/mL to obtain PFU/CFU; the values represent the average of three biological replicates, and error bars represent the standard deviation. A one-way ANOVA was performed with a Tukey’s post-hoc test; the number of phage released per cell in the LB, SGF, and SIF treatments varied significantly from the mitomycin C treatment (p < 0.0005), but not from each other. (B) KP7 and MP13 growth curve. KP7 and MP13 were grown overnight in biological triplicates, OD^600^ was standardized to 1.0, then each strain was sub-cultured 1:100 into LB in 9 technical replicates in a 96-well plate. The plate was incubated in a plate-reader for 15.5 hours at 37 °C with OD^600^ measurements taken every 30 minutes, preceded by a brief shaking interval. Values plotted are the mean of 27 replicates, (excluding 2 outliers for KP7 which did not grow), with the standard deviation shown as error bars. (C) Prophage stability assay; MP13 was grown in LB, 50% LB 50% SGF (29), 50% LB 50% SIF (29), or LB supplemented with 0.5 ng/uL mitomycin C for 24 hours, then sub-cultured into fresh media and grown for another 24 hours. After 48 hours total incubation, cells were plated onto LB agar, then replica plated onto a second LB agar plate spread with KP7 (not lysogenic for Kapi1). The number of non-lysogenic colonies was divided by the total number of colonies to obtain % loss of Kapi1; values represent the average of three biological replicates, and error bars represent the standard deviation. A one-way ANOVA was performed on arcsin-transformed values with a Tukey post-hoc test; the number of Kapi1 lysogens lost in the mitomycin C treatment differed significantly from the LB treatment, and from the SIF treatment (P<0.01).

In order to study the impact of Kapi1 carriage on growth rate, we compared growth of a Kapi1 lysogen, MP13, to that of the non-lysogenized KP7. Kapi1 slightly reduces the growth rate of the host, and results in a lower cell density in stationary phase (Figure 4B). The cell density of lysogens in stationary phase is also more variable than their non-lysogenic counterparts (see error bars in Figure 4B), likely due to spontaneous induction in a subset of the lysogen population (Figure 4A). The time-course of prophage liberation also agrees with the variation in the stationary phase cell density of lysogens, since Kapi1 seems to be induced at the highest levels during stationary phase (data not shown). This is expected, as stationary phase includes stressful cellular conditions such as nutrient limitation and waste product build-up, resulting in the induction of cellular stress responses, stimulating phage release (for a review of stationary phase in Gram-negative bacteria, see (28)).

Since MP1 was recently isolated from the feces of a healthy mouse (11) and is more host-adapted than our standard laboratory strains of *E. coli* such as MG1655 or MC4100, we wondered if Kapi1 might be important to the biology of commensal *E. coli* in the gastrointestinal tract. To investigate the biology of Kapi1 under more physiologically-relevant conditions, we repeated the same prophage liberation experiments in media composed of 50% LB and 50% simulated intestinal fluid (SIF) as well as 50% LB and 50% simulated gastric fluid (SGF) (29). We found lower levels of Kapi1 liberation in SIF when compared to LB (Figure 4A), although the strains grow to nearly identical cell densities. We observed nearly identical levels of Kapi1 liberation in SGF compared to LB (Figure 4A). Prophage stability assays show that upon repeated sub-culturing of a lysogen, the percentage of the population carrying Kapi1 reduces by approximately 10% upon each successive sub-culture in LB, and that the lysogen population is more stable in media composed of 50% LB and 50% SIF (Figure 4C). This result supports our previous finding that the original MP7 isolate lost the Kapi1 prophage before arriving at our lab, likely during handling or passaging.

The prophage liberation and stability assays agree; there is a lower level of Kapi1 induction when lysogens are grown in LB supplemented with SIF compared to LB alone or LB supplemented with SGF, and this low level of induction is likely responsible for the increased stability of the lysogen population in media supplemented with SIF. This result is unexpected because LB is considered a non-stressful standard lab media, yet there is a higher proportion of spontaneous phage induction in this condition, compared to a less rich (and perhaps more challenging) simulated intestinal media. As well, since the liberation rates in SGF were nearly identical to LB, it seems likely that the lower rate of liberation observed in SIF is not simply due to dilution of the rich LB media and may be specific to intestinal conditions. This could indicate that lysogeny with Kapi1 is selected for in intestinal conditions because it provides some advantage to the cell, which may be dispensable under standard lab conditions. Prophages have been shown to provide fitness advantages to their hosts, including resistance to osmotic, oxidative and acid stresses, as well as influencing biofilm formation (30); many of these stresses would be encountered during colonization of the mammalian gastrointestinal tract. Our observations regarding the stability of Kapi1 lysogens in simulated intestinal conditions warrants further investigation into how (or if) bacteriophage influence *E. coli* colonization of the gastrointestinal tract.

### The Kapi1 receptor is lipopolysaccharide O-antigen

Many phage use lipopolysaccharide (LPS) as a receptor for host cell infection, particularly *Podoviridae* (31), including phages HK620 (32) and P22 (18), among others. Based on this knowledge, we hypothesized that Kapi1 may use LPS as its receptor. We isolated spontaneous mutants resistant to Kapi1 by picking survivor colonies from lawns of KP7 overlaid with Kapi1. Since Kapi1 is temperate, survivor colonies were screened for lysogeny by colony PCR with primers spanning the prophage-chromosome junction, and by growing up survivor colonies overnight and spotting their supernatant onto KP7. Two Kapi1-resistant survivor colonies were verified to not be lysogenized by Kapi1, and were chosen for further analysis (KP61, KP62). LPS profiling by SDS-PAGE and silver staining showed that the Kapi1-resistant mutants have severely truncated LPS compared to the WT (Figure 5A). The genomes of KP61 and KP62 were sequenced, and variant calling was performed using WT KP7 as a reference. Both Kapi1-resistant strains contained only one mutation relative to WT; a 1 bp deletion in the *wzy* polymerase causing a frameshift resulting in a truncation (Figure 5B; Figure S3). Mutations in *wzy* result in synthesis of a complete core LPS, but only 1 O-unit is displayed on the cell surface (33), instead of the usual long-chain O-antigens, consistent with what was observed on silver staining. These results indicate that Kapi1 likely binds the KP7 O-antigen as its primary receptor, however it appears that 1 O-unit is insufficient for recognition, as *wzy* mutants are completely resistant to infection (Figure 5C).

**Figure 5.**
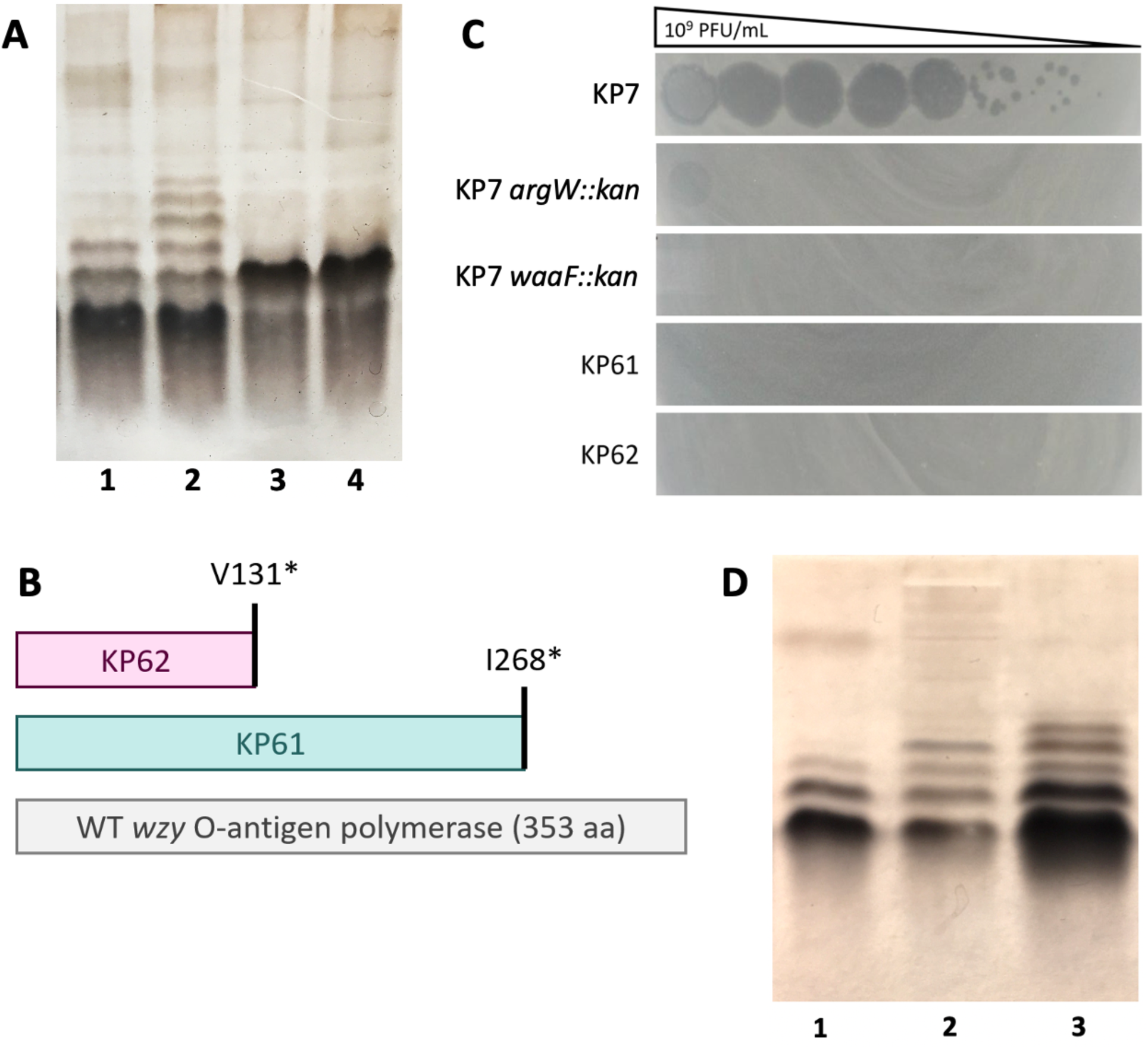
Kapi1 uses the O-antigen as a receptor and modifies its structure. (A) LPS extracted by proteinase K digestion was run on SDS-PAGE and silver stained; lane 1 – WT KP7, lane 2 – MP13 Kap1 lysogen, lane 3 – KP61, lane 4 – KP62. (B) Schematic representing the truncated *wzy* polymerase from KP61 and KP62 compared with the full-length WT protein. (C) Kapi1 phage lysate was serially diluted and spotted onto soft-agar overlays with a variety of mutants in the KP7 background. (D) Phenol-extracted LPS was run on SDS-PAGE and silver stained; lane 1 – KP7, lane 2 – KP7 LPS treated with Kapi1 for 30 minutes, lane 3 – MP13.

To confirm that O-antigen is the receptor for Kapi1, we proceeded to create two LPS mutants; KP7 *waaL::kan* which has complete core, but no O-antigen (34), and KP7 *waaF::kan* which has a severely truncated core structure consisting of lipid A, Kdo_2,_ and heptose (35). Surprisingly, KP7 *waaL::kan* is able to be infected by Kapi1, but at an extremely low efficiency (faint clearing in undiluted spot, Figure 5C), whereas KP7 *waaF::kan* is completely resistant to infection by Kapi1 (Figure 5C). This indicates that in the absence of O-antigen, Kapi1 may be able to recognize a portion of the outer-core structure that is intact in *waaL::kan* but absent in *waaF::kan*, as a secondary receptor; in KP61 and KP62 the single O-unit may obscure this secondary receptor to prevent Kapi1 recognition/binding. Interestingly, *E. coli* W3110 *waaF* mutants have been shown to not produce flagella (36), so this presents another putative secondary receptor for Kapi1. However, no individual plaques can be observed when Kapi1 is spotted on KP7 *waaL::kan* (Figure 5C) so it is also possible that the spot is the result of bactericidal activity, and not a productive phage infection (37). In light of the identification of the O-antigen as the receptor for Kapi1, our previous host-range results make sense; many of the strains we tested lack O-antigen (MC4100, MG1655, BW25113, W3110), and those that do produce O-antigen (E2348/69, J96, DBS100, TJ-LM, TJ-WM, TJ-LR, Nissile 1917) do not appear to have the same O-antigen structure as KP7 on silver-stained SDS-PAGE (data not shown). Although the data represented here is not an extensive screen of all possible serotypes of *E. coli*, it is possible that Kapi1 is specific to one or a few serotypes; identification of the precise region of O-antigen that Kapi1 recognizes as its receptor will help to clarify these results.

### Kapi1 modifies the LPS O-antigen

While performing LPS profiling in the previous experiments, we noticed that the LPS profiles of MP1 and MP13 differed from the LPS profile of KP7 (Figure 5A). We hypothesized that the change in LPS structure was due to lysogenic conversion by Kapi1. Several of the bacteriophage most similar to Kapi1 have been shown to cause seroconversion in their respective hosts. Phages Sf101 and Sf6 both encode O-acetyl transferases to cause seroconversion in their host, *Shigella flexneri* (19, 38); and phage P22 encodes an O-antigen glucosylation cassette (*gtrABC*) to cause seroconversion in its host *Salmonella enterica* serovar Typhimurium (39). We began by searching for CDS in the genome of Kapi1 with homology to known seroconversion proteins. Although none of the CDS in Kapi1 were annotated as possible O-antigen modifying/seroconverting proteins, closer examination revealed limited regions of homology to known seroconversion proteins in a few CDS. Kapi1 hypothetical protein 5 (*hyp5*) has limited homology with acyl- and acetyl-transferases from uncultured Mediterranean phage uvMED (57% identity with 68% query coverage, putative acyltransferase [accession BAR19615.1]; 38% identity with 67% query coverage, putative acetyltransferase [accession B473.1]). Many of the phages infecting *Shigella flexneri* that are closely related to Kapi1 encode seroconversion genes that are located near the *int-att-xis* region of the genome (9), so we included two additional CDS as putative seroconversion proteins due to their location in the genome and lack of homology to any known proteins in the NCBI viral BLASTp database. Kapi1 hypothetical protein 24 (*hyp24*) is encoded between the tail spike protein and the integrase. The only hit in the NCBI viral database is a hypothetical protein, but when the viral filter (taxid: 10239) is removed, this gene shows significant homology with an acyltransferase from *E. coli* (79% identity with 94% query coverage, TPA: acyltransferase [*Escherichia coli*], accession HAH9668903.1). Kapi1 hypothetical protein 25 (*hyp25*) is encoded between the integrase and *attP* site; the only hits in the NCBI database (with or without viral filter) are hypothetical proteins.

We cloned each of these three putative seroconversion proteins into the pTrc99a overexpression vector (40), and introduced them into KP7, then compared the LPS profiles of each overexpression strain, the vector control, and WT KP7 and MP13. Unfortunately, none of the strains overexpressing the three putative seroconversion proteins from Kapi1 had altered LPS profiles (data not shown). Although none of these proteins were found to be individually responsible for seroconversion, we cannot eliminate the possibility that they could be working with other phage-encoded proteins to produce the altered LPS phenotype. There are numerous examples of seroconverting phage that encode entire gene cassettes responsible for seroconversion, including *P. aeruginosa* phage D3 (41) and *Salmonella* phage P22 (39). An alternative approach would be to individually delete putative seroconversion proteins from Kapi1 and assay lysogens for O-antigen structural changes back to the WT form. It is interesting, however, that the genetic basis for O-antigen modification by Kapi1 appears to not be well-conserved, as no strong hits to known O-antigen modifying proteins could be identified. It will be valuable to determine the molecular mechanisms behind Kapi1 phage-mediated O-antigen modification, and whether these mechanisms are indeed novel.

Phages that use the O-antigen as a receptor also commonly modify the O-antigen using the tailspike protein (*tsp*) to facilitate movement to the bacterial outer membrane, where irreversible binding and particle opening can occur (42). To determine if O-antigen degradation via *tsp* was responsible for the altered LPS structure between Kapi1 lysogens and non-lysogens observed on SDS-PAGE (Figure 5A), we set up a mock infection with Kapi1 and purified KP7 LPS and ran these samples alongside the uninfected controls from KP7 and MP13. It appears that treatment of purified KP7 LPS with Kapi1 results in an altered LPS structure (suggesting degradation of O-antigen by Kapi1), but this structure is different from that of either the WT or lysogenic backgrounds (Figure 5D). Therefore, Kapi1 is responsible for alteration of the LPS structure both upon binding O-antigen prior to infection, and later via lysogenic conversion. Kapi1 *tsp* has considerable sequence conservation in the head-binding domain (most similar to phages HK620 and Sf101), but no sequence similarity to any other viral proteins in the NCBI database along the length of the protein. Therefore, it is difficult to predict what type of enzymatic activity the Kapi1 *tsp* may have, and further work is needed to characterize the molecular mechanisms of Kapi1-mediated O-antigen modification through *tsp*.

In conclusion, we have isolated and begun the characterization of a novel bacteriophage infecting commensal *E. coli.* The genome of Kapi1 has been sequenced and annotated, and the prophage integration site in the host genome has been identified. Further, we demonstrate that Kapi1 shows unstable lysogeny, and lysogeny appears to be selected for in intestinal conditions. O-antigen is the Kapi1 receptor and Kapi1 appears to modify the host O-antigen upon initial binding, and later in infection through lysogenic conversion, although the molecular mechanisms are yet to be elucidated. Our findings suggest that Kapi1 lysogeny may confer some advantage during colonization of the intestine and demonstrate that this novel temperate phage alters the structure of LPS, a major determinant of bacterial interaction with the immune system. MP1 and Kapi1 will serve as a good model system to explore what role(s) temperate phages may play in colonization of the gastrointestinal tract by commensal strains of *E. coli*.

## Materials and Methods

### Bacterial Strains and Growth Conditions

Strains MP1, MP7, and MP13 were a generous gift from the Goulian lab (11). A complete list of bacterial strains used in this study can be found in Table S1. Unless otherwise specified, all strains were grown in LB (10 g/L tryptone, 5 g/L NaCl, 5 g/L yeast extract) supplemented with the appropriate antibiotics, at 37 °C and 220 rpm. When plated, cells were grown on LB 1.5% agar supplemented with the appropriate antibiotics and inverted at 37 °C. Antibiotic concentrations used are as follows: ampicillin 100 µg/mL, kanamycin 30 or 50 µg/mL.

### Phage Isolation, Propagation, Host Range, and Transmission Electron Microscopy

Phage were isolated from overnight cultures of MP1 or MP13 by pelleting cells, and filter-sterilizing the supernatant using a 0.45 µm syringe-driven filter. Individual plaques were isolated, propagated, and phage stocks prepared by previously described methods (43), slightly modified. Briefly, the above phage-containing supernatant was mixed with susceptible host strain KP7 1:1, then 3.5 mL soft agar (LB 0.7% agar) was added, the mixture was poured onto solid LB agar plates, and incubated overnight. Individual plaques were picked with a sterile Pasteur pipette and gently resuspended in 500 µL modified suspension media (SM) (50 mM Tris–HCl (pH 7.4), 100 mM NaCl, 10 mM MgSO_4_), this suspension was then amplified using the soft agar overlay technique as above. Plates with near-confluent lysis were used to prepare high-titre stocks by collecting the soft agar layer as follows: SM was poured onto the surface of the plate, and soft agar was gently scraped into a 50 mL Falcon tube using a sterile scoopula, rocked at room temperature for 1 hour and centrifuged to pellet the soft agar. The supernatant was filter-sterilized using a 0.22 µm syringe-driven filter, and stored at 4 °C. Plaque diameter was measured from 10 plaques, using Fiji software (44); the mean and standard deviation are reported.

Phage samples were prepared for transmission electron microscopy (TEM) and imaged by previously described methods using uranyl acetate as a background stain (43) at the University of Alberta Advanced Microscopy Facility. Virion measurements were performed using Fiji software (44) from 44 phage particles; the mean and standard deviation are reported.

The host range for Kapi1 was determined by growing up strains of interest overnight in LB and then 50 µL of overnight culture was added to 3.5 mL soft agar and poured onto a LB plate. Once solidified, Kapi1 lysate was serially diluted and spotted onto each strain. In parallel, a whole plate overlay was prepared as above, using 300 µL undiluted phage lysate and 50 µL overnight culture. The following day, plates were scored for presence or absence of plaques, with KP7 included as a positive control. The whole plate overlays were collected as above, for each strain tested. These “trained” Kapi1 lysates were serially diluted and spotted back onto the same strain to see if Kapi1 host range could be expanded by extended incubation with a particular host, as compared to the first round of spotting.

### Genome Sequencing and PCR

Pure cultures were grown by streaking from glycerol cryo-stocks, then picking single colonies and growing in LB broth overnight. DNA was extracted from overnight cultures using the Lucigen MasterPure Complete DNA and RNA Purification Kit, and the concentration and quality of gDNA was checked using the NanoDrop 2000c. Library preparation and whole genome sequencing was performed by the Microbial Genome Sequencing Centre (MiGS, Pittsburgh PA). Libraries were prepared with the Illumina Nextera kit, and sequenced using the NextSeq 550 platform.

Standard PCR was performed using Taq polymerase (Invitrogen) following the manufacturer’s directions; a single colony was suspended in 20 µL nuclease-free water and 5 µL of this suspension was used per 50 µL reaction. Phage DNA was extracted using a standard phenol/chloroform extraction protocol from the Center for Phage Technology at Texas A&M (45), and 1-5 µL of phage DNA was used per 50 µL reaction. PCR products were checked by running 10 µL on a 2% agarose gel and staining with ethidium bromide. All PCR primers used in this study can be found in Table S2. Sanger sequencing was performed by the Molecular Biology Service Unit at the University of Alberta.

### Genome Assembly, Annotation, and Taxonomy

Paired-end reads of 2×150 bp received from MiGS were uploaded to the public server at usegalaxy.org (46). Galaxy, and all tools there-in, was used for bacterial genome assembly, annotation, and analysis as follows. Illumina adapters and low-quality reads were trimmed using Trim Galore! (47), reads were then checked for quality using FastQC (48) and MultiQC (49). Trimmed reads were *de novo* assembled using Unicycler (50), functioning as a SPAdes (51) optimizer as no long-read data was generated. Quality of assemblies was assessed using Quast (52), and bacterial genomes were then annotated using Prokka (53, 54). SnapGene software (from Insightful Science; available at snapgene.com) and Geneious Prime 2019.2.3 (http://www.geneious.com/) were used for genome visualization, creating genome maps, and designing primers.

The Kapi1 genome was manually annotated using previously described methods (55), slightly modified. This method uses a rigorous approach to score potential coding sequences (CDS) based on various parameters; low-scoring CDS are then discarded, and the remaining CDS are analyzed to determine their correct start codon, again based on a scoring system. Briefly, the prophage genome was run through three auto-annotation programs; GenemarkS (56), Glimmer3 (57) at CPT Phage Galaxy public server (<u>cpt.tamu.edu/galaxy-pub</u>), and Prokka (53) at usegalaxy.org (54). The coding potential for each putative CDS was determined using GenemarkS coding potential graph (56). Putative CDS were then searched against NCBI’s non-redundant protein database using BLASTp (14), and scored based on whether they had significantly similar hits (as determined by the E-value), and whether those hits were known proteins or hypotheticals. Each CDS was scored based on the length of overlap with neighboring CDS, as extremely long overlaps are unlikely, while short overlaps of 1, 4 or 8 bp are more favorable as these suggest organization into an operon. Finally, CDS were scored based on the length of the ORF, where extremely short CDS are penalized the most heavily. Low scoring CDS were discarded prior to start codon identification. Start codons for each CDS were scored in a similar manner, using again the coding potential graph from GenemarkS (56), the number of auto-annotation programs that selected the start codon, sequence similarity matches in NCBI, and length of the ORF. These parameters are listed in order of most important to least and were used to select the most likely start codon for each CDS. CD-Search (58) was also performed for all CDS to assist with functional assignment.

Taxonomic evaluation was performed using vContact2 v9.8 (17) through the CyVerse platform (www.cyverse.org). The analysis was run with the default parameters, using NCBI Bacterial and Archaeal Viral RefSeq V85 (with ICTV + NCBI taxonomy) as the reference database. The resulting network was visualized using CytoScape (59). Duplicated edges were removed from the network (edges represent connections between two nodes, in this case viruses), and only first-neighbors to Kapi1 were kept (nodes that have a direct connection to the Kapi1 node). An edge-weighted, spring-embedded layout was used so that nodes that are more closely related appear closer together spatially, and edges were weighted so that stronger connections (ie. more sequence similarity between two viruses) appear darker and thicker.

### Rates of phage liberation and prophage stability assays

The rate of phage liberation was determined by enumerating CFUs and PFUs in cultures of MP13 Kapi1 lysogens over time. Three colonies of MP13 and three colonies of KP7 (non-lysogenic control) were picked and grown overnight at 37 °C, the next day cultures were adjusted to OD_600_ 1.0 to ensure equal cell numbers, then sub-cultured 1:100 into LB, LB with 0.5 ng/µL mitomycin C, LB mixed 50-50 with simulated intestinal fluid (SIF – 6.8 g KH_2_PO_4_, 1.25 g pancreatin, 3 g bile salts in 1 L dH2O, pH adjusted to 7 (29)), or LB mixed 50-50 with simulated gastric fluid (SGF – 2 g/L NaCl, 3.2 g/L porcine mucosa pepsin, pH adjusted to 3.5 (29)). After 24 hours incubation, an aliquot was taken from each culture, cells were spun down, washed in phosphate-buffered saline (PBS) (137 mM NaCl, 2.7 mM KCl, 10 mM Na_2_HPO_4_, 1.8 mM KH_2_PO_4_), serially diluted, plated onto LB, and grown overnight to enumerate the number of cells in the culture. In parallel, the culture supernatants (containing phage) were serially diluted, spotted onto KP121 soft agar overlays, and incubated overnight at 30 °C to enumerate the number of phage particles in the culture. Phage released per cell was estimated by dividing PFU/mL by CFU/mL. KP121 was used to enumerate phage as this strain has a much lower efficiency of lysogeny with Kapi1 compared to WT KP7, allowing for more accurate enumeration.

Efficiency of lysogeny was determined by infecting KP7 and KP121 with Kapi1 at a multiplicity of infection of 10 at 37 °C for 30 minutes, then washing and plating out surviving cells. The following day, 30 survivor colonies were picked and grown up in liquid culture overnight. Overnight cultures were assayed for lysogeny with Kapi1 in two ways: first, soft-agar overlays were prepared with each survivor culture, and spotted with Kapi1; second, survivor cells were spun down, and the supernatant was spotted onto a soft-agar overlay prepared with KP7. Kapi1 lysogens are resistant to lysis by Kapi1, and their supernatants produce plaques on KP7.

Prophage stability was assayed by serially propagating cultures of MP13 Kapi1 lysogens. Cultures were grown in biological triplicates (three independent colonies) in either LB, LB mixed 50-50 with SGF, LB mixed 50-50 with SIF, or LB with 0.5 ng/µL mitomycin C for 24 hours, then sub-cultured into fresh media and grown for another 24 hours. After a total of 48 hours incubation (1 passage) cells were spun down and washed twice in PBS, serially diluted, plated on LB and grown overnight to get individual colonies. Using velvet squares, colonies were replicated onto a LB plate spread with 50 µL of an overnight culture of KP7 to screen for lysogens (60). This replica plating technique results in two distinct phenotypes: colonies that produce a zone of clearing in the KP7 lawn are scored as lysogens, and colonies without a zone of clearing are scored as non-lysogenic (see Figure S4 for examples and experimental verification of these phenotypes). The relative loss of Kapi1 lysogens in the culture was calculated by dividing the non-lysogen CFU/mL by the total CFU/mL.

### Isolation of Spontaneous Kapi1-resistant mutants

Spontaneous Kapi1-resistant mutants were isolated by spotting Kapi1 onto a soft-agar overlay prepared with KP7. The following day, six colonies that grew within the cleared phage spots were picked and re-streaked onto LB. The following day, these colonies were screened for lysogeny with Kapi1 using PCR with primers that span the phage-genome junction. Two colonies that did not produce phage-genome bands were grown up overnight, then the supernatant was filter-sterilized and spotted onto a soft-agar overlay prepared with KP7. The absence of plaques on KP7 confirmed that these two mutants are not Kapi1 lysogens. To identify which mutations were responsible for the Kapi1-resistant phenotype, each colony was sent for whole genome sequencing at MiGS, as above. SNPs were identified using BreSeq (61) and snippy (62) through Galaxy (46), using the KP7 genome as a reference.

### Generation of Mutants

To create *waaF* mutants, P1 transduction was performed as previously described (63, 64) using the corresponding Keio collection mutant (65) as a donor. The Lambda Red system (66) was used to generate KP7 *waaL::kan*, and KP7 *argW::kan*. Briefly, primers were designed using 50-ntd 5’ extensions homologous to the regions surrounding the gene of interest (H1/H2) with 20-ntd 3’ extensions homologous to the regions flanking the FRT sites and Kan cassette in pKD13 (P1/P2) (65). PCR was performed with Phusion polymerase to amplify the FRT-flanked Kan cassette from pKD13, then these products were purified using the QIAquick PCR Purification Kit and electroporated into KP7 carrying the helper plasmid pKD46. Recovery was performed at 37 °C for 2-3 hours, cells were then plated on LB with 30 µg/mL kanamycin and grown at 37 °C for 24 hours to select for recombinant strains. Kanamycin-resistant colonies were then struck onto LB with 50 µg/mL kanamycin and grown overnight at 37 °C, then patched onto LB with or without ampicillin to screen for loss of pKD46. Kanamycin-resistant and ampicillin-sensitive strains were confirmed by colony PCR using primers designed to span the upstream and downstream junctions between the genome and kanamycin cassette, and primers that spanned the entire disrupted region. PCR products were sequenced at the University of Alberta Molecular Biology Service Unit to confirm the desired mutations. Once sequencing confirmed the correct mutation, P1 transduction was used (as above) to move these mutations into a fresh KP7 background, to avoid the possibility of any off-site mutations acquired during construction. In all cases, strains were confirmed to be non-lysogenic for Kapi1 by PCR and spotting assays, as above.

### Lipopolysaccharide Profiling

The general structure of lipopolysaccharide (LPS) was analyzed by profiling LPS extracts on SDS-PAGE with silver staining. LPS was extracted using a modified proteinase K micro-digestion protocol (67) as follows; bacterial strains of interest were grown overnight, then 1 mL of this culture was washed twice with PBS, resuspended to a final OD_600_ of 2.0 in PBS, and pelleted. Cells were resuspended in 50 µL lysis buffer (2% SDS, 4% 2-mercaptoethanol, 10% glycerol, 1M Tris pH 6.8, bromophenol blue to a deep blue color) and incubated at 95 °C for 10 minutes to lyse cells. Whole cell lysate was cooled to room temperature, then 10 µL 2.5 mg/mL proteinase K (20 mg/mL stock solution was diluted in lysis buffer first) was added and incubated at 56 °C for 1 hour. Standard polyacrylamide gels were prepared with 12% acrylamide (19:1 acrylamide:bisacrylamide) (68), 1-5 µL of proteinase K-treated whole cell lysate (LPS extract) was loaded per well and run at 20 mA constant current in Tris-glycine running buffer (25 mM Tris, 200 mM glycine, 0.1% SDS) until the dye front nearly reached the bottom of the gel. Silver staining was performed as previously described (69), and imaged on a clear petri dish using an iPhone camera.

### Cloning

Putative seroconversion proteins were cloned into pTrc99a (40) using a basic PCR-based cloning method. Coding sequences for the candidate seroconversion proteins were amplified from MP13 colonies using Phusion polymerase; forward primers were designed with an EcoRI restriction site, and reverse primers included a BamHI restriction site. Four 50 µL PCR reactions were pooled for each gene, then cleaned up using the QIAquick PCR Purification Kit. Purified PCR products and empty pTrc99a were double digested with EcoRI and BamHI FastDigest enzymes (Thermo Scientific). Digested vector and inserts were again pooled and cleaned up using the QIAquick PCR Purification Kit. Inserts were ligated into pTrc99a using a 3:1 molar ratio, using T4 DNA ligase (Invitrogen). 8 µL of each ligation was transformed into One Shot TOP10 Chemically Competent *E. coli* (Invitrogen) and were selected on LB plates containing ampicillin. Ampicillin-resistant colonies were screened for inserts by PCR, using primers binding just outside of the multiple cloning site in pTrc99a; colonies with the correct size insert were grown up overnight, the plasmids were isolated using QIAprep Spin Miniprep Kit, then sequenced at the University of Alberta Molecular Biology Service Unit. After verification of the sequences, plasmids were electroporated into KP7. Briefly, electrocompetent cells were prepared by harvesting mid-log phase cells, washing four times in ice-cold 10% glycerol and concentrating cells 500-fold. Electrocompetent cells were combined with 1-5 µL plasmid DNA, electroporated at 2.5 kV in a 0.2 mm cuvette, immediately resuspended in 1 mL LB, recovered for 2-3 hours at 37 °C with shaking, and then plated on LB with ampicillin to select for transformants. To screen candidate seroconversion proteins, KP7 transformants carrying pTrc99a constructs were sub-cultured 1:50 into LB with ampicillin and grown to an OD_600_ of ≥ 0.5, then IPTG was added to a final concentration of 1 mM and cells were grown until an OD_600_ of ≥ 1.0 (20-60 mins of induction) before LPS extraction, SDS-PAGE, and silver staining, as above. All transformants were verified to be non-lysogenic for Kapi1 using the replica plating method described above.

### Lipopolysaccharide Degradation

To determine if Kapi1 is capable of degrading LPS, LPS was extracted from cultures of KP7 (non-lysogenic for Kapi1) and MP13 Kapi1 lysogens using a phenol extraction method adapted from Davis and Goldberg (70), purified KP7 LPS was then incubated with Kapi1 and compared to the untreated controls. The phenol extraction was used in place of proteinase K digestion (as above), as Kapi1 was not viable in the lysis buffer, even after inactivation of proteinase K at high temperatures (data not shown). Briefly, KP7 and MP13 were grown up overnight, cultures were pelleted and washed in PBS, then resuspended in 1.5 mL PBS to give a final OD^600^ of 2.0. Cells were pelleted and resuspended in 200 µL Laemmli buffer (50 mM Tris-HCl pH 6.8, 4% SDS, 10% glycerol, 0.1% bromophenol blue, 5% β-mercaptoethanol), then boiled for 15 minutes to lyse cells. Once cool, DNase I and RNase A were added to cell lysates at 37 °C for 10 minutes. Proteinase K was then added, and lysates were incubated at 55 °C overnight. The following day, 200 µL Tris-saturated phenol was added to lysates, then vortexed for 10 seconds before incubating at 65 °C for 15 minutes. Once cool, samples were centrifuged at 14,000 rpm for 10 min at 4 °C, and the upper phase was transferred to a new tube. 2.5 vol ethanol was added to precipitate LPS, then centrifuged at 15,000 rpm for 20 minutes. Supernatant was discarded, pellets air dried, then resuspended in 50 µL nuclease-free water. Kapi1 was added to KP7 LPS at a “MOI” of 10 (assuming that 1.5 mL of OD^600^ 2.0 culture was concentrated into 50 µL) and incubated at 37 °C without shaking for 30 minutes. Phage-treated and untreated KP7 LPS, along with MP13 untreated LPS were run on SDS-PAGE and silver stained, as above, using 15 µL of LPS extract, as this extraction method produced a lower yield. A portion of the phage-treated samples were serially diluted and spotted onto an KP7 soft-agar overlay to ensure that phage particles remained viable after incubation with LPS (data not shown).

### Data Availability

The genome of Kapi1 can be accessed from NCBI GenBank (accession MT813197).

## Acknowledgments

The authors thank Arlene Oatway from the University of Alberta Advanced Microscopy Facility for assistance with transmission electron microscopy; the University of Alberta Molecular Biology Facility for assistance with Sanger sequencing; the Microbial Genome Sequencing Centre for assistance with whole genome sequencing; Dr. Mark Goulian for critical reading of the manuscript and for strains MP1, MP7, MP13; Dr. Ben Willing for strains TJ-LM, TJ-WM, and TJ-LR; Jaclyn McCutcheon for advice on phage isolation and preliminary characterization; and Dr. Brent Weber for advice on identification of the integration site.

This research was supported by operating grants from The National Sciences and Engineering Research Council (NSERC) and The Canadian Institutes of Health Research (CIHR), and a project grant from the AMR – One Health Consortium, funded by the Major Innovation Fund program of the Ministry of Jobs, Economy and Innovation, Government of Alberta, to TR. KP was supported by an NSERC Alexander Graham Bell Canada Graduate Scholarship - Master’s, Walter H Johns Graduate Fellowship, University of Alberta Science Graduate Scholarship, and Susan Eberlein Graduate Scholarship in Genetics.

## References

1. Tenaillon O, Skurnik D, Picard B, Denamur E. 2010. The population genetics of commensal Escherichia coli. Nat Rev Microbiol 8:207–217.

2. Howard-Varona C, Hargreaves KR, Abedon ST, Sullivan MB. 2017. Lysogeny in nature: Mechanisms, impact and ecology of temperate phages. ISME J 11:1511–1520.

3. Touchon M, Bernheim A, Rocha EPC. 2016. Genetic and life-history traits associated with the distribution of prophages in bacteria. ISME J 10:2744–2754.

4. Kim MS, Bae JW. 2018. Lysogeny is prevalent and widely distributed in the murine gut microbiota. ISME J 12:1127–1141.

5. Hsu BB, Gibson TE, Yeliseyev V, Liu Q, Lyon L, Bry L, Silver PA, Gerber GK. 2019. Dynamic Modulation of the Gut Microbiota and Metabolome by Bacteriophages in a Mouse Model. Cell Host Microbe 25:803–814.e5.

6. Frazão N, Sousa A, Lässig M, Gordo I. 2019. Horizontal gene transfer overrides mutation in Escherichia coli colonizing the mammalian gut. Proc Natl Acad Sci U S A 116:17906– 17915.

7. Davies E V., Winstanley C, Fothergill JL, James CE. 2016. The role of temperate bacteriophages in bacterial infection. FEMS Microbiol Lett.

8. Van Belleghem JD, Dąbrowska K, Vaneechoutte M, Barr JJ, Bollyky PL. 2019. Interactions between bacteriophage, bacteria, and the mammalian immune system. Viruses 11.

9. Allison GE, Verma NK. 2000. Serotype-converting bacteriophages and O-antigen modification in Shigella flexneri. Trends Microbiol 8:17–22.

10. Guest RL, Rutherford ST, Silhavy TJ. 2020. Border Control : Regulating LPS Biogenesis. Trends Microbiol 1–12.

11. Lasaro M, Liu Z, Bishar R, Kelly K, Chattopadhyay S, Paul S, Sokurenko E, Zhu J, Goulian M. 2014. Escherichia coli isolate for studying colonization of the mouse intestine and its application to two-component signaling knockouts. J Bacteriol 196:1723–1732.

12. Arndt D, Grant JR, Marcu A, Sajed T, Pon A, Liang Y, Wishart DS. 2016. PHASTER: a better, faster version of the PHAST phage search tool. Nucleic Acids Res 44:W16–W21.

13. Zhou Y, Liang Y, Lynch KH, Dennis JJ, Wishart DS. 2011. PHAST: A Fast Phage Search Tool. Nucleic Acids Res 39:347–352.

14. Altschul SF, Gish W, Miller W, Myers EW, Lipman DJ. 1990. Basic local alignment search tool. J Mol Biol 215:403–410.

15. Adriaenssens EM, Rodney Brister J. 2017. How to name and classify your phage: An informal guide. Viruses 9:1–9.

16. International Committee on Taxonomy of Viruses. 2020. Master Species Lists. Retrieved from https://talk.ictvonline.org/files/master-species-lists/.

17. Bin Jang H, Bolduc B, Zablocki O, Kuhn JH, Roux S, Adriaenssens EM, Brister JR, Kropinski AM, Krupovic M, Lavigne R, Turner D, Sullivan MB. 2019. Taxonomic assignment of uncultivated prokaryotic virus genomes is enabled by gene-sharing networks. Nat Biotechnol 37:632–639.

18. Neal BL, Brown PK, Reeves PR. 1993. Use of Salmonella phage P22 for transduction in Escherichia coli. J Bacteriol 175:7115–7118.

19. Jakhetia R, Marri A, Ståhle J, Widmalm G, Verma NK. 2014. Serotype-conversion in Shigella flexneri: Identification of a novel bacteriophage, Sf101, from a serotype 7a strain. BMC Genomics 15.

20. Clark AJ, Inwood W, Cloutier T, Dhillon TS. 2001. Nucleotide sequence of coliphage HK620 and the evolution of lambdoid phages. J Mol Biol 311:657–679.

21. Nakata K, Koh MM, Tsuchido T, Matsumura Y. 2010. All genomic mutations in the antimicrobial surfactant-resistant mutant, Escherichia coli OW66, are involved in cell resistance to surfactant. Appl Microbiol Biotechnol 87:1895–1905.

22. McFall E, Heincz M. 1983. Identification and control of synthesis of the dsdC activator protein. J Bacteriol 153:872–877.

23. Moritz RL, Welch RA. 2006. The Escherichia coli argW-dsdCXA genetic island is highly variable, and E. coli K1 strains commonly possess two copies of dsdCXA. J Clin Microbiol 44:4038–4048.

24. Reiter W-D, Palm P, Yeats S. 1989. Transfer RNA genes frequently serve as integration sites for prokaryotic genetic elements. Nucleic Acids Res 17.

25. Casjens S, Winn-Stapley DA, Gilcrease EB, Morona R, Kühlewein C, Chua JEH, Manning PA, Inwood W, Clark AJ. 2004. The chromosome of Shigella flexneri bacteriophage Sf6: Complete nucleotide sequence, genetic mosaicism, and DNA packaging. J Mol Biol 339:379–394.

26. Panis G, Méjean V, Ansaldi M. 2007. Control and regulation of KplE1 prophage site-specific recombination: A new recombination module analyzed. J Biol Chem 282:21798– 21809.

27. Barreiro V, Haggard-Ljungquist E. 1992. Attachment sites for bacteriophage P2 on the Escherichia coli chromosome: DNA sequences, localization on the physical map, and detection of a P2-like remnant in E. coli K-12 derivatives. J Bacteriol 174:4086–4093.

28. Navarro Llorens JM, Tormo A, Martínez-García E. 2010. Stationary phase in gram-negative bacteria. FEMS Microbiol Rev. Blackwell Publishing Ltd.

29. Millette M, Nguyen A, Amine KM, Lacroix M. 2013. Gastrointestinal survival of bacteria in commercial probiotic products. Int J Probiotics Prebiotics 8:149–156.

30. Wang X, Kim Y, Ma Q, Hong SH, Pokusaeva K, Sturino JM, Wood TK. 2010. Cryptic prophages help bacteria cope with adverse environments. Nat Commun 1:147.

31. Silva JB, Storms Z, Sauvageau D. 2016. Host receptors for bacteriophage adsorption. FEMS Microbiol Lett 363:1–11.

32. Dhillon TS, Poon APW, Chan D, Clark AJ. 1998. General transducing phages like Salmonella phage P22 isolated using a smooth strain of Escherichia coli as host. FEMS Microbiol Lett 161:129–133.

33. Samuel G, Reeves P. 2003. Biosynthesis of O-antigens: Genes and pathways involved in nucleotide sugar precursor synthesis and O-antigen assembly. Carbohydr Res 338:2503– 2519.

34. Heinrichs DE, Monteiro MA, Perry MB, Whitfield C. 1998. The assembly system for the lipopolysaccharide R2 core-type of Escherichia coli is a hybrid of those found in Escherichia coli K-12 and Salmonella enterica. Structure and function of the R2 WaaK and WaaL homologs. J Biol Chem 273:8849–8859.

35. Gronow S, Brabetz W, Brade H. 2000. Comparative functional characterization in vitro of heptosyltransferase I (WaaC) and II (WaaF) from Escherichia coli. Eur J Biochem 267:6602–6611.

36. Wang Z, Wang J, Ren G, Li Y, Wang X. 2016. Deletion of the genes waaC, waaF, or waaG in Escherichia coli W3110 disables the flagella biosynthesis. J Basic Microbiol 56:1021–1035.

37. Abedon ST. 2018. Detection of Bacteriophages: Phage Plaques, p. 1–32. *In* Bacteriophages.

38. Verma NK, Brandt JM, Verma DJ, Lindberg AA. 1991. Molecular characterization of the O-acetyl transferase gene of converting bacteriophage SF6 that adds group antigen 6 to Shigella flexneri. Mol Microbiol 5:71–75.

39. Vander Byl C, Kropinski AM. 2000. Sequence of the genome of Salmonella bacteriophage P22. J Bacteriol 182:6472–6481.

40. Amann E, Ochs B, Abel KJ. 1988. Tightly regulated tac promoter vectors useful for the expression of unfused and fused proteins in Escherichia coli. Gene 69:301–315.

41. Newton GJ, Daniels C, Burrows LL, Kropinski AM, Clarke AJ, Lam JS. 2001. Three-component-mediated serotype conversion in Pseudomonas aeruginosa by bacteriophage D3. Mol Microbiol 39:1237–1247.

42. Broeker NK, Barbirz S. 2017. Not a barrier but a key: How bacteriophages exploit host’s O-antigen as an essential receptor to initiate infection. Mol Microbiol 105:353–357.

43. Peters DL, Mccutcheon JG, Stothard P, Dennis JJ. 2019. Novel Stenotrophomonas maltophilia temperate phage DLP4 is capable of lysogenic conversion. BMC Genomics 20.

44. Schindelin J, Arganda-Carreras I, Frise E, Kaynig V, Longair M, Pietzsch T, Preibisch S, Rueden C, Saalfeld S, Schmid B, Tinevez JY, White DJ, Hartenstein V, Eliceiri K, Tomancak P, Cardona A. 2012. Fiji: An open-source platform for biological-image analysis. Nat Methods 9:676–682.

45. Center for Phage Technology. 2018. Protocol for Phage DNA Extraction with Phenol:Chloroform. Retrieved from https://cpt.tamu.edu/phage-links/phage-protocols.

46. Afgan E, Baker D, Batut B, Van Den Beek M, Bouvier D, Ech M, Chilton J, Clements D, Coraor N, Grüning BA, Guerler A, Hillman-Jackson J, Hiltemann S, Jalili V, Rasche H, Soranzo N, Goecks J, Taylor J, Nekrutenko A, Blankenberg D. 2018. The Galaxy platform for accessible, reproducible and collaborative biomedical analyses: 2018 update. Nucleic Acids Res 46:W537–W544.

47. Babraham Bioinformatics. 2012. Trim Galore! A wrapper tool around Cutadapt and FastQC to consistently apply quality and adapter trimming to FastQ files. Retrieved from https://www.bioinformatics.babraham.ac.uk/projects/trim_galore/.

48. Babraham Bioinformatics. 2010. FastQC, A quality control tool for high throughput sequence data. Retrieved from http://www.bioinformatics.babraham.ac.uk/projects/fastqc/.

49. Ewels P, Magnusson M, Lundin S, Käller M. 2016. MultiQC: Summarize analysis results for multiple tools and samples in a single report. Bioinformatics 32:3047–3048.

50. Wick RR, Judd LM, Gorrie CL, Holt KE. 2017. Unicycler: Resolving bacterial genome assemblies from short and long sequencing reads. PLoS Comput Biol 13:1–16.

51. Bankevich A, Nurk S, Antipov D, Gurevich AA, Dvorkin M, Kulikov AS, Lesin VM, Nikolenko SI, Pham S, Prjibelski AD, Pyshkin A V., Sirotkin A V., Vyahhi N, Tesler G, Alekseyev MA, Pevzner PA. 2012. SPAdes: A new genome assembly algorithm and its applications to single-cell sequencing. J Comput Biol 19:455–477.

52. Mikheenko A, Prjibelski A, Saveliev V, Antipov D, Gurevich A. 2018. Versatile genome assembly evaluation with QUAST-LG. Bioinformatics 34:i142–i150.

53. Seemann T. 2014. Prokka: Rapid prokaryotic genome annotation. Bioinformatics 30:2068–2069.

54. Cuccuru G, Orsini M, Pinna A, Sbardellati A, Soranzo N, Travaglione A, Uva P, Zanetti G, Fotia G. 2014. Orione, a web-based framework for NGS analysis in microbiology. Bioinformatics 30:1928–1929.

55. Salisbury A, Tsourkas PK. 2019. A method for improving the accuracy and efficiency of bacteriophage genome annotation. Int J Mol Sci 20.

56. Besemer J. 2001. GeneMarkS: a self-training method for prediction of gene starts in microbial genomes. Implications for finding sequence motifs in regulatory regions. Nucleic Acids Res 29:2607–2618.

57. Delcher A, Bratke K, Powers E, Salzberg S. 2007. Identifying bacterial genes and endosymbiont DNA with Glimmer. Bioinformatics 23:673–679.

58. Marchler-Bauer A, Lu S, Anderson JB, Chitsaz F, Derbyshire MK, DeWeese-Scott C, Fong JH, Geer LY, Geer RC, Gonzales NR, Gwadz M, Hurwitz DI, Jackson JD, Ke Z, Lanczycki CJ, Lu F, Marchler GH, Mullokandov M, Omelchenko M V., Robertson CL, Song JS, Thanki N, Yamashita RA, Zhang D, Zhang N, Zheng C, Bryant SH. 2011. CDD: A Conserved Domain Database for the functional annotation of proteins. Nucleic Acids Res 39:225–229.

59. Shannon P, Markiel A, Ozier O, Baliga NS, Wang JT, Ramage D, Amin N, Schwikowski B, Ideker T. 2003. Cytoscape: A Software Environment for Integrated Models. Genome Res 13:2498–2504.

60. Lederberg EM, Lederberg J. 1953. Genetic Studies of Lysogenicity in Escherichia coli. Genetics 38:51–64.

61. Deatherage DE, Barrick JE. 2014. Identification of mutations in laboratory evolved microbes from next-generation sequencing data using breseq. Methods Mol Biol https://doi.org/10.1007/978-1-4939-0554-6_12.

62. Seemann T. 2015. snippy: fast bacterial variant calling from NGS reads. Retrieved from https://github.com/tseemann/snippy.

63. Sambrook J, Russell DW. 2001. Molecular cloning: a laboratory manual 3rd edition. Coldspring Harbour Laboratory Press, UK.

64. Silhavy TJ, Berman ML, Enquist LW. 1984. Experiments with gene fusions. Cold Spring Harb Lab.

65. Baba T, Ara T, Hasegawa M, Takai Y, Okumura Y, Baba M, Datsenko KA, Tomita M, Wanner BL, Mori H. 2006. Construction of Escherichia coli K-12 in-frame, single-gene knockout mutants: The Keio collection. Mol Syst Biol https://doi.org/10.1038/msb4100050.

66. Datsenko KA, Wanner BL. 2000. One-step inactivation of chromosomal genes in Escherichia coli K-12 using PCR products. Proc Natl Acad Sci U S A 97:6640–6645.

67. Hitchcock PJ. 1984. Analyses of gonococcal lipopolysaccharide in whole-cell lysates by sodium dodecyl sulfate-polyacrylamide gel electrophoresis: Stable association of lipopolysaccharide with the major outer membrane protein (protein I) of Neisseria gonorrhoeae. Infect Immun 46:202–212.

68. Kulikov EE, Golomidova AK, Prokhorov NS, Ivanov PA, Letarov A V. 2019. High-throughput LPS profiling as a tool for revealing of bacteriophage infection strategies. Sci Rep 9.

69. Tsai C, Frasch C. 1982. Silver Stain for Detecting Lipopolysaccharides Polyacrylamide Gels. Anal Biochem 19:115–119.

70. Davis MR, Goldberg JB. 2012. Purification and visualization of lipopolysaccharide from gram-negative bacteria by hot aqueous-phenol extraction. J Vis Exp 1.

71. Wintersinger JA, Wasmuth JD. 2015. Kablammo: An interactive, web-based BLAST results visualizer. Bioinformatics 31:1305–1306.

72. Guyer MS, Reed RR, Steitz JA, Low KB. 1981. Identification of a sex-factor-affinity site in E. coli as gamma delta. Cold Spring Harb Symp Quant Biol 45:135–140.

73. Ju T, Shoblak Y, Gao Y, Yang K, Fouhse J, Finlay B, Wing So Y, Stothard P, Willing BP. 2017. Initial Gut Microbial Composition as a Key Factor Driving Host Response to Antibiotic Treatment, as Exemplified by the Presence or Absence of Commensal Escherichia coli. Appl Environ Microbiol 83.

74. Casadaban MJ. 1976. Transposition and fusion of the lac genes to selected promoters in Escherichia coli using bacteriophage lambda and Mu. J Mol Biol 104:541–555.

75. Bachmann BJ. 1972. Pedigrees of some mutant strains of Escherichia coli K-12. Bacteriol Rev 36:525–557.

76. Hull RA, Gill RE, Hsu P, Minshew BH, Falkow S. 1981. Construction and expression of recombinant plasmids encoding type 1 or D-mannose-resistant pili from a urinary tract infection Escherichia coli isolate. Infect Immun 33:933–938.

77. Iguchi A, Thomson NR, Ogura Y, Saunders D, Ooka T, Henderson IR, Harris D, Asadulghani M, Kurokawa K, Dean P, Kenny B, Quail MA, Thurston S, Dougan G, Hayashi T, Parkhill J, Frankel G. 2009. Complete genome sequence and comparative genome analysis of enteropathogenic Escherichia coli O127:H6 strain E2348/69. J Bacteriol 91:347–354.

78. Nissile A. 1918. Die antagonistische Behandlung chronischer Darmstörungen mit Colibakterien. Med Klin 29–30.

79. Schauer DB, Falkow S. 1993. The eae gene of Citrobacter freundii biotype 4280 is necessary for colonization in transmissible murine colonic hyperplasia. Infect Immun 61:4654–4661.

